# High titer methyl ketone production with tailored *Pseudomonas taiwanensis* VLB120

**DOI:** 10.1101/2020.06.02.125906

**Authors:** Salome C. Nies, Tobias B. Alter, Sophia Nölting, Susanne Thiery, An N. T. Phan, Noud Drummen, Jay D. Keasling, Lars M. Blank, Birgitta E. Ebert

## Abstract

Methyl ketones present a group of highly reduced platform chemicals industrially produced from petroleum-derived hydrocarbons. They find applications in the fragrance, flavor, pharmacological, and agrochemical industries, and are further discussed as biodiesel blends. In recent years, intense research has been carried out to achieve sustainable production of these molecules by re-arranging the fatty acid metabolism of various microbes. One challenge in the development of a highly productive microbe is the high demand for reducing power. Here, we engineered *Pseudomonas taiwanensis* VLB120 for methyl ketone production as this microbe has been shown to sustain exceptionally high NAD(P)H regeneration rates. The implementation of published strategies resulted in 2.1 g L_aq_^-1^ methyl ketones in fed-batch fermentation. We further increased the production by eliminating competing reactions suggested by metabolic analyses. These efforts resulted in the production of 9.8 g L_aq_^-1^ methyl ketones (corresponding to 69.3 g L_org_^-1^ in the *in situ* extraction phase) at 53 % of the maximum theoretical yield. This represents a 4-fold improvement in product titer compared to the initial production strain and the highest titer of recombinantly produced methyl ketones reported to date. Accordingly, this study underlines the high potential of *P. taiwanensis* VLB120 to produce methyl ketones and emphasizes model-driven metabolic engineering to rationalize and accelerate strain optimization efforts.

## 1. Introduction

The conversion of renewable feedstocks or waste streams by microbial fatty acid biosynthesis into commodity oleochemicals and transportation fuels represents a sustainable alternative to petroleum or plant-oil based processes. The range of molecules derived from fatty acids includes free fatty acids, esters, alcohols, aldehydes, ketones, alkanes, olefins, or waxes (Kim et al., 2019; Xu et al., 2016). These molecules find applications as (advanced) biofuels, flavors, fragrances, detergents, lubricants, surfactants, or plastic monomers (Buist, 2010; Lennen and Pfleger, 2013; Tee et al., 2014) and are derived from either fatty acyl Coenzyme A (CoA) or fatty acyl-acyl carrier protein thioesters. The essentiality of these metabolites for lipid biosynthesis in most living organisms (Janssen and Steinbüchel, 2014; López-Lara and Geiger, 2010) guarantees constitutive and substantial production, and successful examples show the possibility of rewiring microbial metabolism for the synthesis of fatty acid-derived products at high titers (Kim et al., 2019; Xu et al., 2016).

Fatty acid-derived methyl ketones are garnering attention for use as flavors, fragrances, insecticides or insect repellents, solvents as well as for their promising fuel properties (Goh et al., 2012; Kimps et al., 2011; Longo, 2006). The cetane number describes the ignition behavior of diesel fuels (Dahmen and Marquardt, 2015) and has been determined for a mixture of medium-chain length methyl ketones (C11 and C13) to be around 58 (Goh et al., 2012). While this is already in the diesel range, the cetane number can be further increased to 80 to 90 by chemical modification of methyl ketones to dioxolanes (Harrison and Harvey, 2018). Microbes do not naturally synthesize methyl ketones, but recombinant production of medium-chain length congeners has been achieved in several species (Dong et al., 2019; Goh et al., 2012; Hanko et al., 2018; Müller et al., 2013; Park et al., 2012). However, most reported titers are in the mg L^-1^ range, and lower gram-scale production has only been reported for an intensively engineered *E. coli* in a 200-h fed-batch cultivation (Goh et al., 2018).

Besides overexpression of the product biosynthesis pathway, a typical strategy to boost the synthesis of fatty acid derivatives is engineering the supply of the precursors acetyl-CoA and malonyl-CoA. This can be achieved, e.g., by overexpressing acetyl-CoA carboxylase, catalyzing the first committed step of fatty acid synthesis (Davis et al., 2000) or deletion of acetyl-CoA consuming pathways. Indeed, abolishing acetyl-CoA conversion to acetate was crucial for improving methyl ketone production in *E. coli*. (Goh et al., 2014). Secretion of fermentative byproducts is not only a loss of carbon but also of electrons. The latter is vital since fatty acid synthesis has a high demand for NADPH, and the supply of electrons via reduced redox coenzymes can limit the production rate. The formation of reducing equivalents can be shifted to NADPH by overexpression of a transhydrogenase (Sanchez et al., 2006), NADH kinase (Lee et al., 2009) or by changing the cofactor dependency of key enzymes. While such interventions positively impacted methyl ketone production (Goh et al., 2018), they do not increase the overall reductive power of the cell. Redox cofactor availability, therefore, remains a nontrivial challenge.

In this study, we decided on utilizing *P. taiwanensis* VLB120, an obligate aerobic bacterium as host for methyl ketone production. Pseudomonads are gaining increasing interest in industrial biotechnology due to their fast growth, solvent tolerance, and versatile metabolism (Kohler et al., 2013; Poblete-Castro et al., 2012a). Relevant for the envisaged application, Pseudomonads produce no fermentative byproducts and can sustain an exceptionally high redox cofactor regeneration rate depending on the metabolic demand (Blank et al., 2008; Ebert et al., 2011; Rühl et al., 2009). The natural ability of *P. taiwanensis* VLB120 to utilize glycerol, xylose (Kohler et al., 2015), and aromatic compounds found in lignocellulosic biomass hydrolysate (Panke et al., 1998; Poblete-Castro et al., 2012a) renders the strain interesting for biorefinery approaches.

We argue hence that *P. taiwanensis* VLB120 offers excellent traits for the production of methyl ketones. To prove its potential, we first implemented the methyl ketone synthesis pathway previously established in *E. coli* (Goh et al., 2012; Goh et al., 2014) and applied a metabolic model-supported engineering approach that boosted methyl ketone titer to 9.8 g L_aq_^-1^ and a yield of 53 % of the theoretical maximum.

## 2. Material and Methods

### 2.1. Strains, media, and culture conditions

Table 1 lists the bacterial strains used in this study. Strains were propagated in Lysogeny Broth (LB) containing 10 g L^-1^ peptone, 5 g L^-1^ sodium chloride, and 5 g L^-1^ yeast extract (Sambrook, 1982). Cetrimide agar (Sigma-Aldrich, St. Louis, MO, USA) was used after the mating procedures to select for *Pseudomonas*. Solid LB was prepared by adding 1.5 % (w/v) agar to the liquid medium. For plasmid maintenance, the gene deletion procedure, and genomic integrations, antibiotics were added to the medium as required. Gentamycin, kanamycin, and ampicillin were used at concentrations of 25 mg L^-1^, 50 mg L^-1^, and 100 mg L^-1^, respectively. *E. coli* was grown at 37 °C, *Pseudomonas* at 30 °C on a horizontal rotary shaker with a throw of 50 mm and a frequency of 300 rpm (Infors, Bottmingen, Switzerland). The chemicals used in this work were obtained from Carl Roth (Karlsruhe, Germany), Sigma-Aldrich (St. Louis, MO, USA), or Merck (Darmstadt, Germany) unless stated otherwise.

**Table 1.**
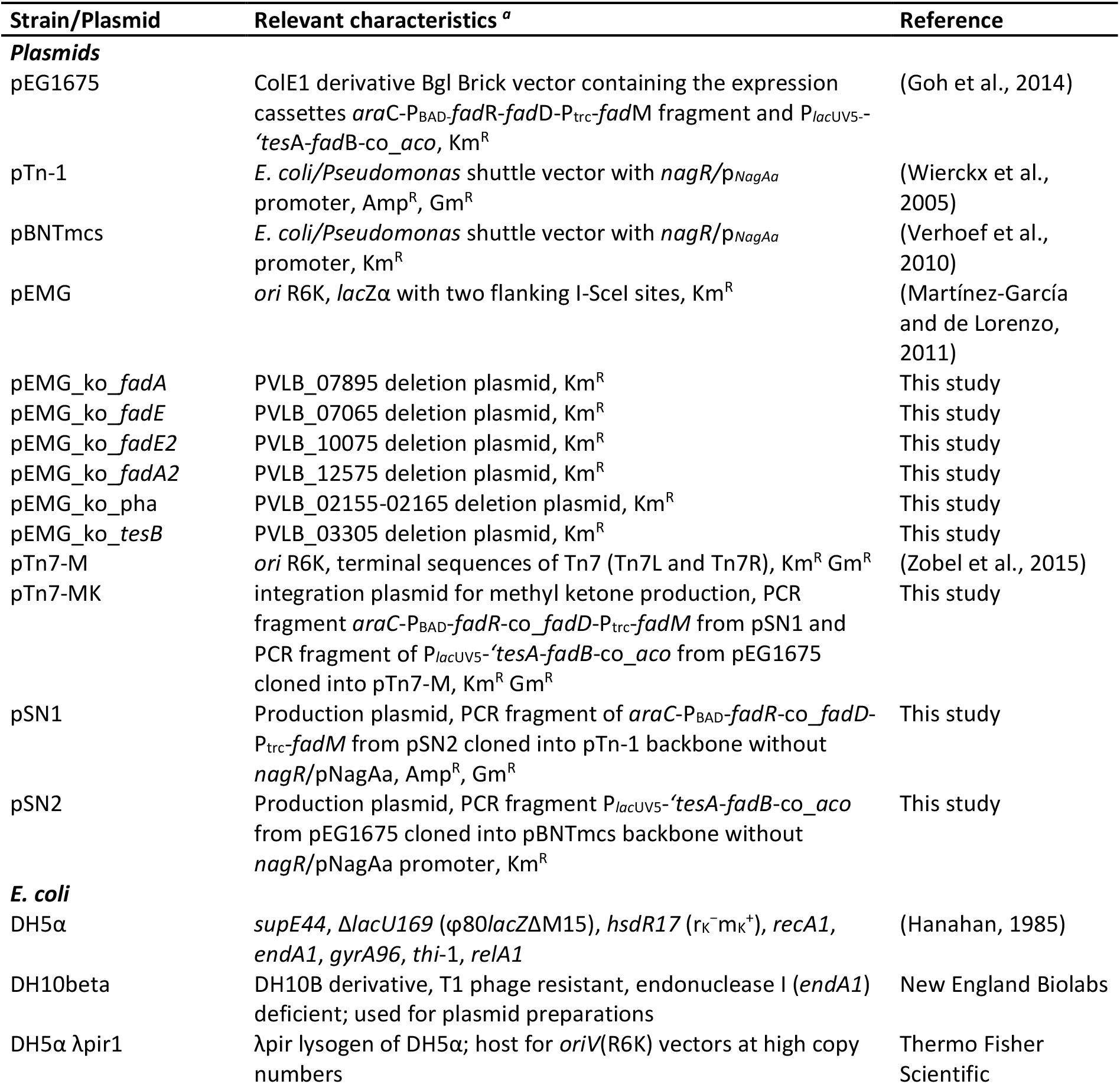
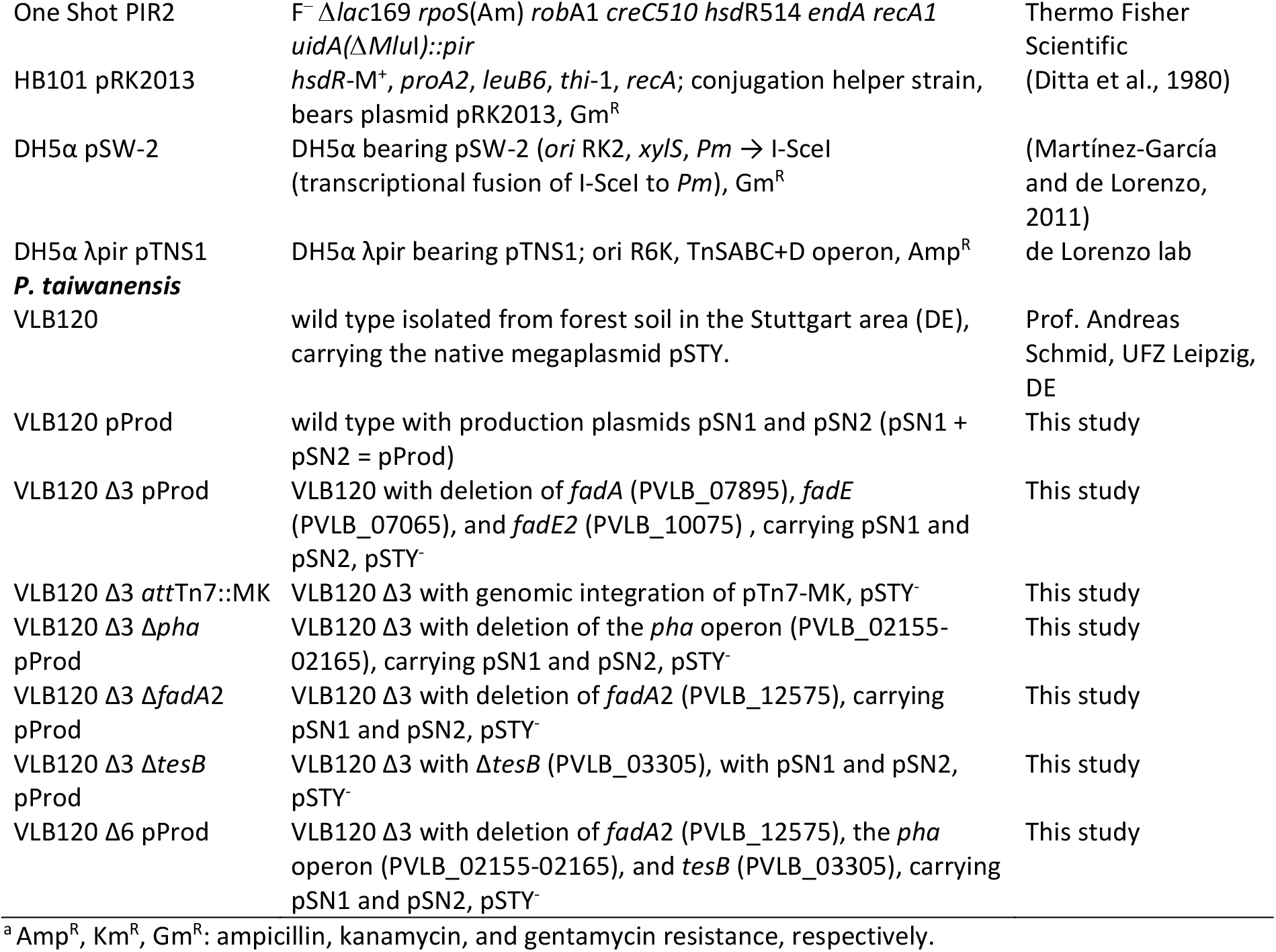
Bacterial strains and plasmids used in this study.

Methyl ketone production by the engineered *Pseudomonas* was characterized by cultivating the strains in 500-mL flasks with 10 % (v/v) mineral salt medium (Hartmans et al., 1989) containing 7.76 g L^-1^ K_2_HPO_4_, 3.26 g L^-1^ NaH_2_PO_4_, 2 g L^-1^ (NH_4_)_2_SO_4_, 0.1 g L^-1^ MgCl_2_·6H_2_O, 10 mg L^-1^ EDTA, 2 mg L^-1^ ZnSO_4_·7 H_2_O, 1 mg L^-1^ CaCl_2_·2H_2_O, 5 mg L^-1^ FeSO_4_·7 H_2_O, 0.2 mg L^-1^ Na_2_MoO_4_·2H_2_O, 0.2 mg L^-1^ CuSO_4_·5 H_2_O, 0.4 mg L^-1^ CoCl_2_·6 H_2_O, 1 mg L^-1^ MnCl_2_·2 H_2_O supplemented with 25 mM or 50 mM glucose unless stated otherwise. For continuous *in situ* extraction of methyl ketones, 12.5 mL *n*-decane was added as an overlay. If not indicated otherwise samples for analytical measurements were taken after 24 h and 48 h. Before inoculating the preculture, LB agar plates (with appropriate antibiotics) were streaked from frozen glycerol stocks and were incubated overnight at 30 °C. A single colony was used to inoculate 5 mL of mineral salt medium containing antibiotics, supplemented with 25 mM glucose in a 15-mL glass tube. The preculture was transferred twice before inoculating the main culture. The main cultures were inoculated from liquid precultures to an approximate optical density at 600 nm (OD_600nm_) of 0.1. Gene expression was induced after 5-7 h with the addition of 2 mM IPTG and 1 mM arabinose for plasmid-bearing strains. Arabinose and IPTG were added to a final concentration of 1 mM for gene expression in strains with a genome integrated methyl ketone pathway. All experiments were performed in biological duplicates unless stated otherwise.

### 2.2. Plasmid construction and transformation

Primers were ordered as unmodified DNA oligonucleotides from Eurofins Genomics (Ebersberg, Germany) and are listed in Supplementary Table S1. The Monarch Plasmid Miniprep Kit (New England Biolabs, Ipswich, MA, USA) was used to isolate plasmids, which were either digested with restriction enzymes purchased from New England Biolabs (Ipswich, MA, USA) or were used as a template for PCR using Q5 High-Fidelity Polymerase (New England Biolabs, Ipswich, MA, USA). The plasmid pEG1675 developed by Goh et al. (Goh et al., 2014) for methyl ketone production in *E. coli* served as a template for the amplification of two genetic cassettes required for methyl ketone overproduction. All genes, except co_*aco*, originate from *E. coli; co_fadD* had been codon-optimized for high expression in *E. coli*. ‘*tesA* encodes a truncated variant of the thioesterase TesA that expresses in the cytosol; co_*aco* designates the acyl-CoA oxidase encoding gene Mlut_11700 from *Micrococcus luteus* codon-optimized for *E. coli*. The plasmids pSN1 and pSN2 were introduced into chemically competent *E. coli* DH10beta (New England Biolabs, Ipswich, MA, USA). Transformants were screened by colony PCR using OneTaq 2x Master Mix (New England Biolabs, Ipswich, MA, USA). Production plasmids were isolated and analyzed by PCR and Sanger sequencing (Eurofins Genomics, Germany).

For gene deletions, upstream (TS1) and downstream (TS2) regions with a length of 400-800 bp flanking the specific target gene were amplified using Q5 High-Fidelity Polymerase (New England Biolabs, Ipswich, MA, USA). Genomic DNA of *P. taiwanensis* VLB120 was used as the template for the PCR and was isolated using the High Pure PCR Template Preparation Kit (Hoffmann-La-Roche, Basel, Switzerland). Plasmids were constructed by Gibson assembly using the NEBuilder Hifi DNA Assembly Master Mix (New England Biolabs, Ipswich, MA, USA). pEMG plasmids were transformed into competent *E. coli* DH5α λpir1 via electroporation (Choi et al., 2006), and the correct assembly was confirmed by colony PCR. The cell material was lysed in alkaline polyethylene glycol for enhanced colony PCR efficiency as described elsewhere (Chomczynski and Rymaszewski, 2006).

### 2.3. Generation of deletion strains and genomic integration

Targeted gene deletions were performed using the I-SceI-based system developed by Martinez-Garcia and de Lorenzo (Martínez-García and de Lorenzo, 2011) and as described previously (Wynands et al., 2018). Briefly, the conjugational transfer of the mobilizable knockout plasmid from *E. coli* DH5α λpir1 to *Pseudomonas* was performed via patch mating overnight (Wynands et al., 2018). After conjugation, the pSW-2 plasmid encoding the I-SceI-endonuclease was conjugated into *Pseudomonas* co-integrates. The induction of I-SceI expression with 3-methyl benzoate was omitted as the basal expression level was sufficient. Kanamycin-sensitive clones were directly isolated, positive clones were cured of pSW-2, and re-streaked several times. The gene deletion was confirmed by colony PCR as described above and by Sanger sequencing.

Genomic integration into the *att*Tn7 of *Pseudomonas* was carried out as described previously (Zobel et al., 2015). Briefly, patch mating was carried out with *E. coli* pRK2013 (helper strain), *E. coli* DH5α-λpir pTnS-1 (strain leading transposase), *E. coli* DH5α One Shot PIR 2 pTn7-MK and VLB120 Δ3. After 48 h of conjugation at 30 °C, cell material was selected on cetrimide agar containing gentamycin. Colonies were screened for positive insertion with colony PCR as described above.

### 2.4. Analysis of cell growth and sugar metabolism during aqueous organic two-phase cultivations

Samples taken from aqueous organic two-phase cultivations with *n*-decane were treated as follows. First, the different phases were separated by centrifugation for 5 min at 17,000 x g, and the interphase was marked. The organic phase was removed for gas chromatography (GC) analysis, whereas the remaining aqueous phase was filtered through a 0.2 μm membrane filter and stored at −20 °C until further high-performance liquid chromatography (HPLC) analysis. As *n*-decane would perturb OD_600nm_ measurements, residual *n*-decane was carefully removed from the inner tube wall with a cellulose tissue. The cell pellet was resuspended in 0.9 % (w/v) NaCl to a final volume equaling the initial aqueous phase. The OD_600nm_ was determined in technical triplicates using an Ultrospec 10 spectrophotometer (GE Healthcare, Chicago, IL, USA).

Glucose and gluconate concentrations were determined by HPLC. The analysis was performed using a Beckman System Gold 126 Solvent Module equipped with a System Gold 166 UV-detector (Beckman Coulter) and a Smartline RI detector 2300 (KNAUER Wissenschaftliche Geräte, Berlin, Germany). Analytes were separated on the organic resin column Metab AAC (ISERA, Düren, Germany) eluted with 5 mM H2SO4 at an isocratic flow of 0.6 mL min^-1^ at 40 °C for 40 min. Glucose and gluconate were analyzed using the RI detector, whereas gluconate was determined with the UV detector at a wavelength of 210 nm.

### 2.5. Methyl ketone analysis with gas chromatography

Quantification of methyl ketones was performed on a Trace GC Ultra (Thermo Scientific, Waltham, MA, USA) equipped with a flame ionization detector (FID). Analyte separation was achieved with a polar ZB-WAX column (30 m length, 0.25 mm inner diameter, 0.25 μm film thickness, Zebron, Phenomenex, UK) and a constant helium flow of 2 mL min^-1^. The initial oven temperature of 80 °C was held for 2.5 min, increased to 250 °C at 20 °C min^-1^ and held constant for 10 min. The injection volume was 1 μL, and the injector temperature was set to 250 °C. The split ratio was adjusted depending on the analyte concentration. External standard quantification with 2-undecanone (Alfa Aesar, Ward Hill, MA, USA), 2-tridecanone (Alfa Aesar, Ward Hill, MA, USA), 2-pentadecanone, and 2-heptadecanone was performed. No authentic standards were available for the monounsaturated methyl ketones (C11:1, C13:1, C15:1, and C17:1). Those congeners were first identified by GC-mass spectrometry (MS) analysis using a TRACE GC Ultra coupled to the triple quadrupole TSQ 8000 (Thermo Fisher Scientific) and identified by comparing the mass spectra with published data (Goh et al., 2012) (Supplementary Figure S1). The tuning and calibration of the mass spectrometer were done prior to analysis. A 1 μL aliquot of the sample was injected into a VF-5ms capillary column (30 m × 250 μm i.d., 0.25 μm film thickness) with a 10 m EZ-Guard column (Agilent). The injection temperature was 250 °C, and the helium carrier gas flow was set to 1 mL/min. The column temperature was held at 80°C for 2 min, increased by 80 °C min^-1^ to 120 °C, increased by 15 °C min^-1^ to 250 °C and then increased by 100 °C min^-1^ to 325 °C, at which it was held constant for 3 min. The transfer line and ion source temperatures were 280 °C and 300 °C, respectively. Ions were generated by electron ionization at 70 eV.

Saturated methyl ketones were quantified with a calibration curve recorded using authentic standards. We assumed the same response factor for the monounsaturated methyl ketones as for their unsaturated counterparts and quantified the concentration using the respective calibration curve. Methyl ketone titers were related to the aqueous phase, as reported previously (Hanko et al., 2018). For batch fermentations, the aqueous concentration *c_aq_* at time point *t_i_* (*c_aq,t_i__*) was calculated using equation 1:

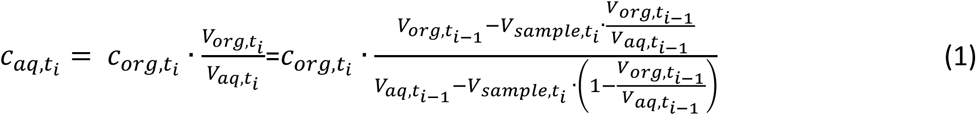

with *c_org, t_i__* being the experimentally determined methyl ketone concentration, and *V_org,t_i__, V*_*org,t*_*i*_−1_ and *V_aq, t_i__, V_aq,t_i__* the volume of the organic, respectively the aqueous phase at time point *t_i_* or *t*_*i*–1_. For fed-batch cultivations, the change in the fermentation volume due to the aqueous medium feed *F_feed_* needed to be considered. For this fermentation mode, the aqueous concentration was calculated using equation 2.

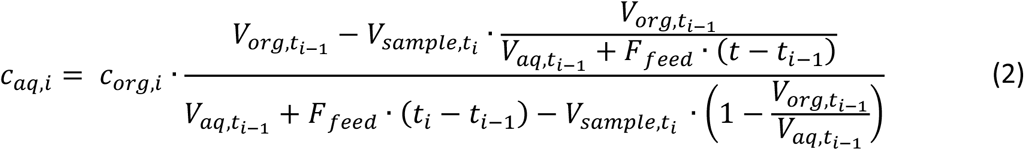

### 2.6. Bioreactor cultivations for methyl ketone production

Bioreactor cultivations were performed in 1.3-L fermenters (BioFlow120, Eppendorf, Germany) with a working volume of 0.5 L mineral salt medium (0.4 L mineral salt medium in fed-batch), supplemented with 2 % (w/v) glucose monohydrate and 100 mL *n*-decane (17 % (v/v)). The temperature was maintained at 30 °C; the pH was controlled at a value of 7 with 2 M KOH and 25 % NH_4_OH in batch and fed-batch cultivations, respectively, and 4 M H_2_SO_4_. The dissolved oxygen (DO) level was kept above 30 % using a stirring cascade. After 6-7 h inoculation, the gene expression was induced with 2 mM IPTG and 1 mM arabinose. For the fed-batch cultivation, the DO signal was used to start the feeding pump, when DO signal exceeded 70 %, the feeding pump started with 5 mL h^-1^, with a feeding medium of 585 mM glucose, 7.76 g L^-1^ K_2_HPO_4_, 3.26 g L^-1^ NaH_2_PO_4_, 2 g L^-1^ (NH_4_)_2_SO_4_, 0.3 g L^-1^ MgCl_2_·6H_2_O, 30 mg L^-1^ EDTA, 6 mg L^-1^ ZnSO_4_·7 H_2_O, 3 mg L^-1^ CaCl_2_·2H_2_O, 15 mg L^-1^ FeSO_4_·7 H_2_O, 0.6 mg L^-1^ Na_2_MoO_4_·2H_2_O, 0.6 mg L^-1^ CuSO_4_·5 H_2_O, 1.2 mg L^-1^ CoCl_2_·6 H_2_O, 3 mg L^-1^ MnCl_2_·2 H_2_O, 2 mM IPTG, 1 mM arabinose, and appropriate antibiotics (gentamycin and kanamycin). The culture broth was sampled regularly for OD_600nm_ determination, extracellular metabolite analysis, and methyl ketone quantification.

### 2.7. Model-guided strain-design

For the model-guided strain design, the genome-scale metabolic model *i*JN1411 of *Pseudomonas putida* KT2440 (Nogales et al., 2017) was employed and manually adapted to match the enzymatic repertoire of *P. taiwanensis* VLB120 (Supplementary file S1). *In silico* synthesis of methyl ketone precursors was enabled by extending the model with a thioesterase III reaction (*fadM*) converting β-ketoacyl-CoA to the respective β-keto acid for each methyl ketone congeners (C11-C17). Additional spontaneous reactions modeled the cleavage of CO2 from the β-keto acid yielding the specific methyl ketones, which are transported to the extracellular space by additional passive transport reactions. The exchange of all considered methyl ketones was pooled in a single reaction with fixed stoichiometric coefficients for each congener. The coefficients were calculated based on the final molar concentration ratios of the most dominant methyl ketones produced by the VLB120 Δ3 pProd (cf. Results, section 3.3). The thioesterase encoded by *fadA* was additionally deleted to fully resemble VLB120 Δ3 pProd. In the following, we refer to *i*JN1411_MK for this final metabolic reconstruction. For the prediction of genetic intervention strategies that enhance methyl ketones production, an optimization algorithm based on a genetic algorithm was applied (Alter et al., 2018). The Supplementary File S2 lists the applied optimization parameters. The genetic algorithm utilizes the principles of natural selection to evolve a random, initial population of solutions towards optimality. Each solution comprises a set of gene deletions that relate to specific reaction deletions in a stoichiometric metabolic model by its inherited gene-protein-reaction relations. The impact of a set of gene deletions on production was evaluated by using the yield Y_GM_ of methyl ketones on glucose as the fitness function. The yield was calculated by Y_GM_ = ν_MK_ / ν_GLC_, where ν_MK_ and ν_GLC_ are the flux of the methyl ketones exchange reaction and the glucose uptake rate, respectively. Both ν_MK_ and ν_GLC_ are variables in the flux distributions computed by the Minimization of Metabolites Balances (MiMBl) algorithm (Brochado et al., 2012). MiMBl calculates mutant flux distributions by minimizing the distance to a reference or wild-type flux distribution with respect to the metabolite balance vector, which comprises the sums of absolute flux rates of all reactions producing or consuming a particular metabolite. We derived a reference flux distribution from the phenotypic flux data of the cultivation of VLB120 Δ3 pProd (cf. Results, section 3.3) using a parsimonious flux balance analysis (pFBA) approach (Lewis et al., 2010). Since data from the exponential growth phase were used to compute the reference steady-state flux distribution, the reference MK yield is significantly lower as compared to the yield at the end of the batch cultivation. During the evolution of a population to near-optimality, genetic algorithms generate a multitude of solutions, thus simultaneously condense the full solution space to a subset of the most beneficial gene deletions. While the ultimate best set of genetic interventions is most promising for methyl ketone overproduction, alternative solutions with a similar fitness may also be valuable candidates for application. Moreover, information about the significance of gene deletions in generating an overproducing phenotype is desirable for deciding on the order by which to apply single genetic interventions. The first step to assist in this practical issue was to extract solutions generated by the genetic algorithm with an increase in methyl ketone yield of at least 50 % compared to the VLB120 Δ3 pProd strain. All gene deletion targets were ranked according to the number of occurrences within these solutions.

## 3. Results and discussion

### 3.1. Metabolic engineering of *P. taiwanensis* VLB120 for methyl ketone production

The ability of *Pseudomonas* to produce methyl ketones was initially assessed following a strategy previously used in *E. coli* (Goh et al., 2012; Goh et al., 2014). This approach included the truncation of the β-oxidation cycle at the 3-ketothiolase (*fadA* deletion), the overexpression of a type III thioesterase (*fadM*) to drain 3-keto-acyl CoAs from the β-oxidation cycle, and ramping up fatty acid synthesis and the subsequent oxidation by overexpression of corresponding genes (Figure 1A).

**Figure 1.**
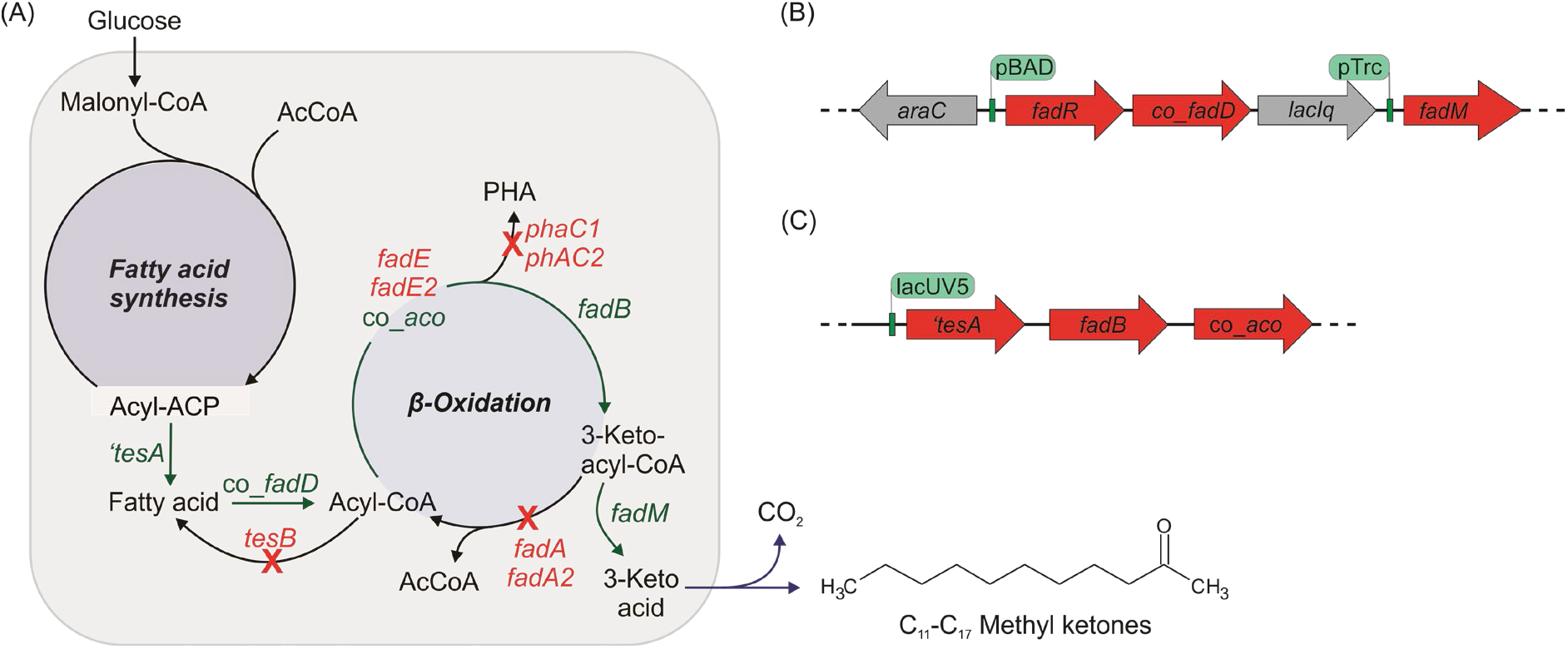
(A) Fatty acid biosynthesis and degradation pathways in *P. taiwanensis* engineered for methyl ketone production. Gene names in green indicate overexpression; those in red represent gene deletion targets. (B/C) Schematic illustration of genetic constructs used for recombinant methyl ketone production in *P. taiwanensis*. The expression cassettes (B) and (C) were either expressed from the plasmids pSN1 (B) and pSN2 (C) or as a combined construct integrated into the genome as a single copy. See text for further details.

For the overexpression of the product synthesis genes, two plasmids were constructed. Plasmid pSN1 (Figure 1B) carried the fatty acid-responsive transcription factor *fadR, fadD*, encoding an acyl-CoA synthetase, and *fadM* encoding a thioesterase III in the pTn1 backbone (Nijkamp et al., 2005; Wierckx et al., 2005). The expression of these genes was under the control of either the inducible pBAD or pTrc promoter. The truncated *‘tesA* gene, encoding a cytosolic thioesterase I, *fadB*, a subunit of the fatty acid oxidation complex with hydratase activity, and a codon-optimized version of the acyl-CoA oxidase gene Mlut_11700 from *Micrococcus luteus* (co_*aco*) were inserted into the pBNTmcs backbone (Verhoef et al., 2010) and expressed under the control of the inducible *lacUV5* promoter (plasmid pSN2, Figure 1C). The co-transformation of the wild type *P. taiwanensis* VLB120 with the two plasmids pSN1 and pSN2 resulted in VLB120 pProd, with pProd designating the two production plasmids pSN1 and pSN2.

The two thioesterases ‘TesA and FadM are characterized by a broad substrate specificity and act on molecules with carbon chain lengths between C12-C16 (‘TesA) and C12-C18 (FadM), respectively (Goh et al., 2012; Nie et al., 2008; Steen et al., 2010). They were chosen to gear the methyl ketones synthesis towards congeners with 11 to 17 carbon atoms for potential use as diesel blend. This distribution was confirmed for the proof-of-concept strain VLB120 pProd, which produced in 24 h 0.6 mg L_aq_^-1^ total methyl ketones in a mineral salt medium and the presence of a second organic layer of *n*-decane (Figure 2). The observed distribution of methyl ketone congeners reflects the substrate specificities of the overexpressed acyl-CoA thioesterases ‘TesA and FadM. The difference to the natural fatty acid profile of *P. taiwanensis* VLB120, which is dominated by C16:0, C16:1, and C18:1 fatty acids (Rühl et al., 2012) indicates that the expressed thioesterase ‘TesA efficiently drains shorter chain length molecules from the fatty acid biosynthesis pathway.

**Figure 2.**
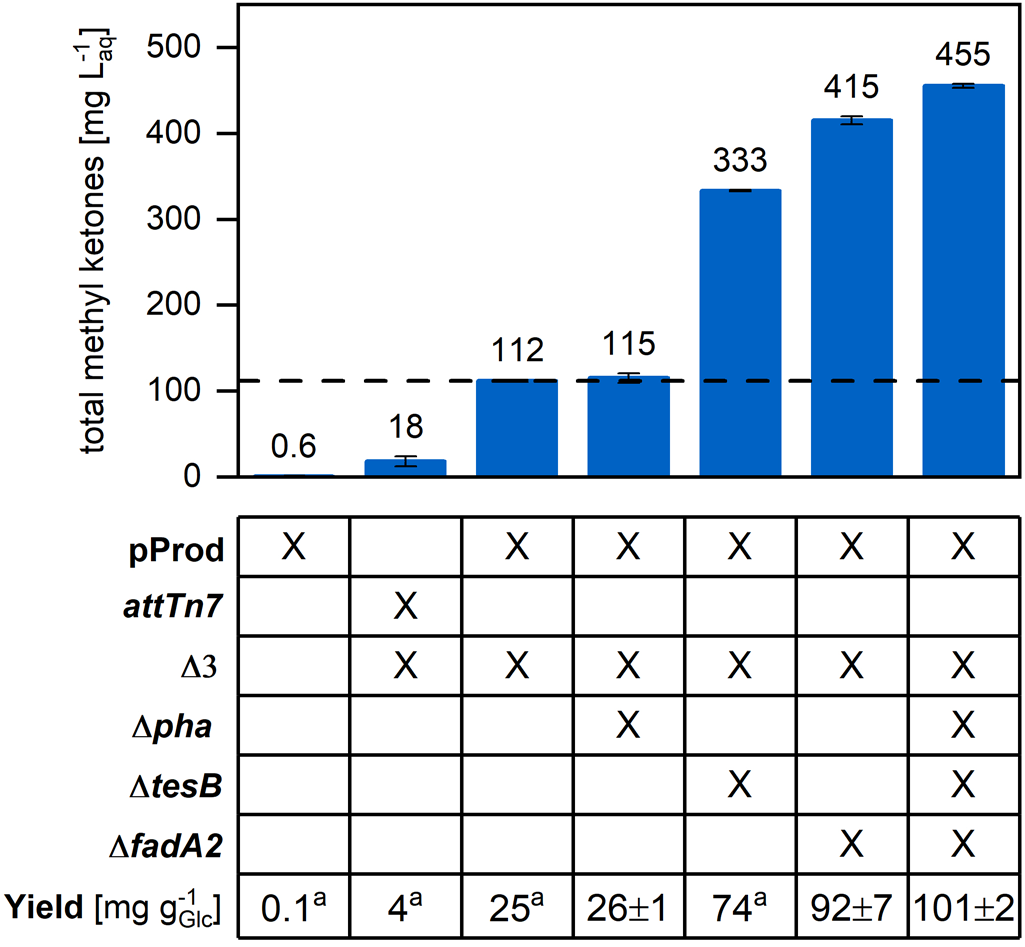
Methyl ketone production in metabolically engineered *P. taiwanensis* VLB120 strains. *att*Tn7 refers to the strain with genome integrated production pathway genes; Δ3 summarizes the deletions of*fadA, fadE*, and *fadE2*; Δ*pha* comprises the deletion of the *pha* operon (polyhydroxyalkanoate synthases and depolymerase). The dotted line highlights the titer of the VLB120 Δ3 pProd strain. All strains were cultivated in mineral salt medium supplemented with 25 mM glucose and 20 % (v/v) *n*-decane for in-situ extraction. Methyl ketone measurements were taken after complete carbon source depletion, i.e., after 24 h or 48 h. Errors represent the standard deviation of data of duplicate experiments; ^a^standard deviation of less than one.

We further modified the genetic background of *P. taiwanensis* VLB120 by deleting the thioesterase encoding gene *fadA* (Goh et al., 2012). This intervention results in a truncation of the β-oxidation cycle, which should enforce the flux towards β-keto acids and their spontaneous decarboxylation to methyl ketones in *fadM* expressing strains (Goh et al., 2012). Since we expressed a heterologous, soluble acyl-CoA oxidase (co_*aco*), we deleted the redundant, native acyl-CoA dehydrogenase encoding genes PVLB_07065 (*fadE*) and PVLB_10075 (here referred to as *fadE2*) in the engineered strain. The resulting triple knockout strain is referred to as VLB120 Δ3.

We used this basis strain to test the effect of different pathway gene expression levels by either expressing the pathway genes from the multi-copy number plasmids pSN1 and pSN2 (VLB120 Δ3 pProd) or a single-copy genome-integrated cassette (VLB120 Δ3 *att*Tn7::MK). While the latter strain will express the pathway genes at a lower level, it might profit from a reduced metabolic burden associated with plasmid replication. To this end, the two fragments (Figure 1B&C) were amplified from the pSN1 and pSN2 plasmids, cloned into the pTn7-M vector, and integrated into the single, genomic *att*Tn7 site using a mini-Tn7 transposon-based method (Zobel et al., 2015).

### 3.2. Characterization of methyl ketone production by plasmid-based and single-copy genome integrated pathway expression

We aimed for a balanced expression of the pathway genes to maximize productivity and therefore determined the optimal inducer concentration for the plasmid-based and chromosomally integrated cassette. With the two-plasmid based approach, the highest titers were achieved with 1 mM arabinose and 2 mM IPTG, whereas 1 mM arabinose and 1 mM IPTG was optimal for the expression of the genome integrated single-copy methyl ketone pathway (Supplementary Figure S2).

The deletion of*fadA* significantly boosted methyl ketone production in these strains. In a 24-h shake flask experiment, VLB120 Δ3 pProd reached 112 mg L_aq_^-1^ total methyl ketones (187-fold compared to VLB120 pProd) with a yield of 24.9 mg g_GLC_^-1^ (Figure 2). As with its predecessor strain, monounsaturated and saturated methyl ketones with a chain length between C11 and C17 were detected.

The replication of the two large production plasmids exerted a high metabolic burden on VLB120 Δ3 pProd evident from a substantial growth defect. This burden was reduced in VLB120 Δ3 *att*Tn7::MK, which grew faster (specific growth rate μ = 0.61 h^-1^ vs 0.44 h^-1^) and reached a higher maximal biomass concentration (OD_600nm_ of ~7 vs ~4). Despite the growth disadvantage, plasmid-based expression of the pathway genes in VLB120 Δ3 pProd outperformed methyl ketone production in VLB120 Δ3 *att*Tn7::MK, which accumulated a total methyl ketone titer of only 18 mg L_aq_^-1^. For this reason, we continued all further work with VLB120 Δ3 pProd.

### 3.3. Performance of *P. taiwanensis* VLB120 Δ3 pProd in bioreactor cultivations

The potential of the strain VLB120 Δ3 pProd was further assessed in pH- and DO-controlled batch and fed-batch fermentations. Under batch conditions, 540 mg L_aq_^-1^ methyl ketones were produced with a productivity of 11 mg L_aq_^-1^ h^-1^ (Figure 3A). The dominant methyl ketones were the saturated C11 and C13 and the monounsaturated C13:1 and C15:1 congeners.

**Figure 3.**
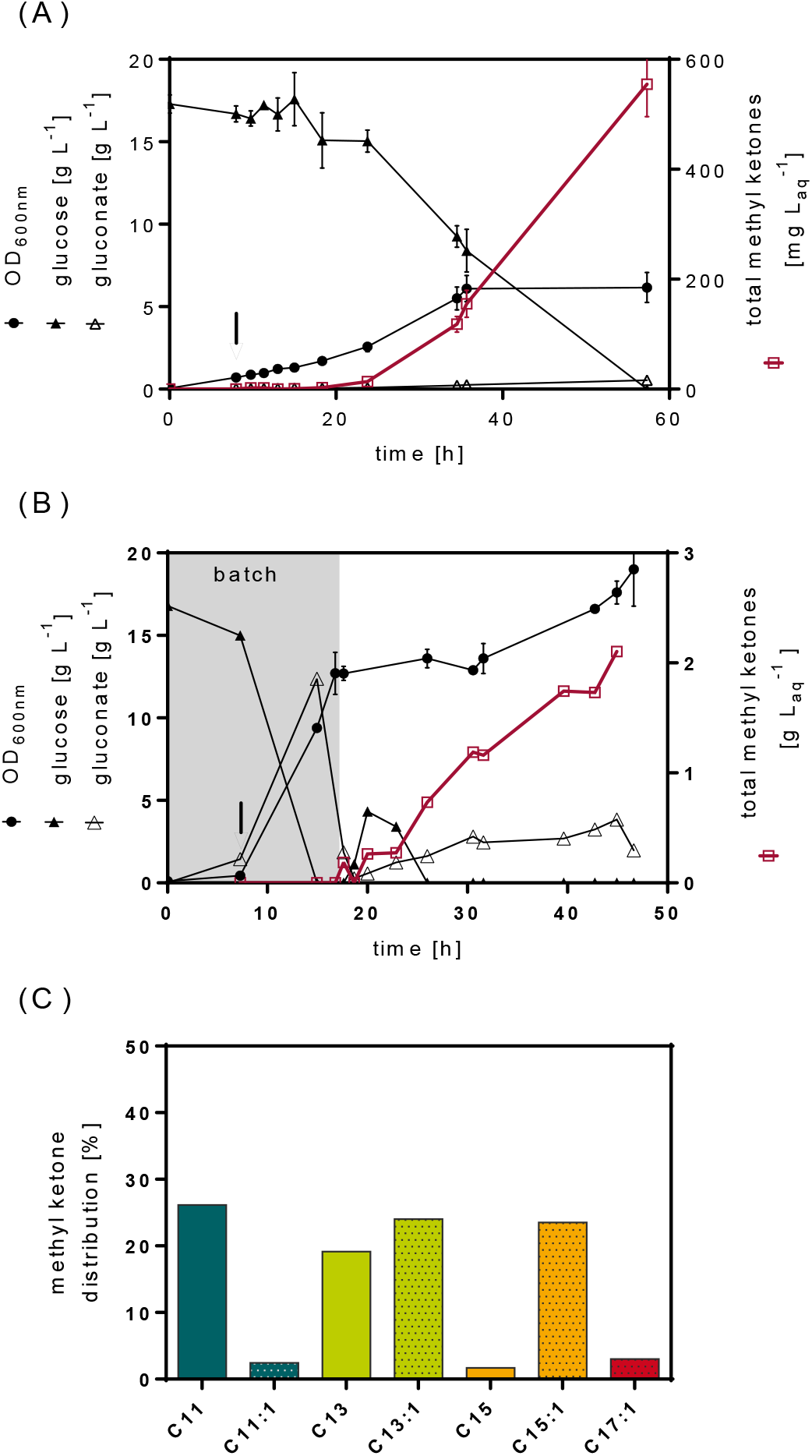
Performance of *P. taiwanensis* VLB120 Δ3 pProd in (A) batch and (B) fed-batch cultivation. Both cultivations were done in mineral salt medium with an initial concentration of 17 % (v/v) *n*-decane as second organic phase. The time point of induction is indicated by a vertical arrow. The total methyl ketones concentration is the sum of all detected saturated and unsaturated congeners (C11, C11:1, C13, C13:1, C15, C15:1, C17:1). (C) Methyl ketone distribution at the endpoint of the fed-batch fermentation.

Significant methyl ketone production was only observed in stationary phase. The cessation of growth occurred upon nitrogen depletion, which was the limiting nutrient in the medium used while carbon was supplied in excess. Additional nitrogen supply via ammonium hydroxide titration for pH control was low as the strain produced only minor amounts of gluconate (Figure 3A).

To gain higher product titers, we extended the production phase by fed-batch fermentation and aimed for a similar growth limitation. Differences in growth and carbon source consumption between the initial batch experiment (Figure 3A) and the batch phase of the fed-batch cultivation (Figure 3B) are explained by an improved pre-cultivation procedure. After the end of the batch phase, glucose (585 mM) in mineral salt medium was fed continuously at a rate of 5 mL h^-1^. This feed regimen results in a significant carbon excess (C:N ratio of 39:1) and potentially nitrogen-limited growth known to upregulate the fatty acid metabolism (Poblete-Castro et al., 2012b) and hence favorable for methyl ketone synthesis. During the fed-batch phase, the methyl ketone concentration increased linearly to a final titer of 2.1 g L_aq_^-1^ (Figure 3B), corresponding to a yield of 50 mg g_GLc_^-1^ (Table 2). The methyl ketone distribution did not change during the feed phase (Figure 3C).

**Table 2.** Summary of key performance metrics of the engineered *P. taiwanensis* VLB120 strains in batch and fed-batch fermentations.

| | VLB120 $\Delta 3$ pProd | | VLB120 $\Delta 3\Delta tesB$ pProd | | VLB120 $\Delta 6$ pProd |
| --- | --- | --- | --- | --- | --- |
|  | batch*** | fed-batch* | batch** | fed-batch** | fed-batch* |
| total MK [g L<sub>aq</sub><sup>-1</sup>] | 0.6 ± 0.1 | 2.1 | 1.3 ± 0.2 | 6.0 ± 0.1 | 9.8 |
| Productivity [mg L<sub>aq</sub><sup>-1</sup> h<sup>-1</sup>] | 11 ± 1 | 56 | 30 ± 5 | 82 ± 18 | 132 |
| yield [mg g<sub>GLC</sub><sup>-1</sup>] | 32 ± 4 | 50 | 76 ± 17 | 117 ± 4 | 171 |

\*\*\* n = 3; \*\* n = 2; \* n = 1; for n > 1, mean values and standard deviations are shown; MK, methyl ketones; GLC, glucose; aq, aqueous phase; the volumetric productivity was determined for the time period after induction of heterologous gene expression.

### 3.4 High-titer methyl ketone production after full truncation of the β-oxidation cycle

Further genome analysis revealed that *P. taiwanensis* VLB120 possesses three isozymes of the acetyl-CoA acyltransferase FadA encoded by *fadA* (PVLB_07895), PVLB_12575 (referred to as *fadA2*), and PVLB_26537. The gene PVLB_26537 is located on *P. taiwanensis* VLB120’s megaplasmid, which the strain had lost during the gene knock-out procedure. We deleted the remaining *fadA2* in *P. taiwanensis* VLB120 Δ3 to fully truncate the β-oxidation cycle and co-transformed the strain with the two production plasmids. In shake flask experiments, *P. taiwanensis* VLB120 Δ3 Δ*fadA*2 pProd produced 415 mg L_aq_^-1^ total methyl ketones, an almost 4-fold increase in methyl ketone titer (Figure 2).

### 3.5 Metabolic modeling predicts genetic engineering targets for improved methyl ketone production

To assess the potential for further advancement of methyl ketone production, we determined the theoretical maximum product yield using a metabolic model of *P. taiwanensis* VLB120 extended with the engineered methyl ketone synthesis pathway (*i*JN1411_MK, cf. Material and Methods section 2.7). With glucose as the sole carbon source and the experimentally determined distribution of methyl ketone congeners, the maximal theoretical yield computes to 325 mg g_GLc_^-1^. The yield of 92 ±7 mg g_GLc_^-1^ achieved in a shake-flask cultivation of VLB120 Δ3 *ΔfadA2* pProd (Figure 2) thus corresponds to about 30 % of the maximum. The redox coenzyme demand required to achieve the maximal methyl ketone yield was computed to 89.6 mmol_NAD(P)H_ gMK^-1^, which is more than double the demand for biomass synthesis (40.6 mmol_NAD(P)H_ g_CDW_^-1^) (Supplementary Figure S3). The highest reported glucose uptake rate for *P. taiwanensis* VLB120 of 10 mmol g_CDW_^-1^ h^-1^ (Rühl et al., 2009) translates into a maximal product synthesis rate of 0.6 g_MK_ g_CDW_^-1^ h^-1^ and a NAD(P)H turnover of 52 mmol g_CDW_^-1^ h^-1^. While this is about 2-fold the rate under standard growth conditions (at a specific growth rate of 0.68 h^-1^ (Nies et al., 2020)), this demand does not challenge the redox cofactor regeneration capacity of *P. taiwanensis* VLB120 reported to reach 149 mmol g_CDW_^-1^ h^-1^ (Rühl et al., 2009). These *in silico* analyses indicate a substantial room for improvement of methyl ketone synthesis in *P. taiwanensis* VLB120 by metabolic engineering.

To develop VLB120 Δ3 Δ*fadA*2 pProd towards more efficient methyl ketone production, we predicted the most favorable strain designs with three to five simultaneous gene deletions using a genetic optimization algorithm and the *i*JN1411_MK reconstruction. The quadruple and quintuple mutants with the highest predicted fitness both showed an increase in methyl ketone yield of about 8-fold compared to the reference, which in these simulations was VLB120 Δ3 pProd. None of the triple gene deletion designs met our criteria for a minimal production enhancement of 50 % and were therefore not further assessed. For intervention sets with four and five gene deletions, the genes *phaC1* and *tesB* were present in 81 % to 100 % of the fittest strain designs and presented the highest observed occurrences (Figure 4). Since neither of these targets had been experimentally tested at this point, we decided to integrate *phaC1* and *tesB* deletions in our engineering efforts.

**Figure 4.**
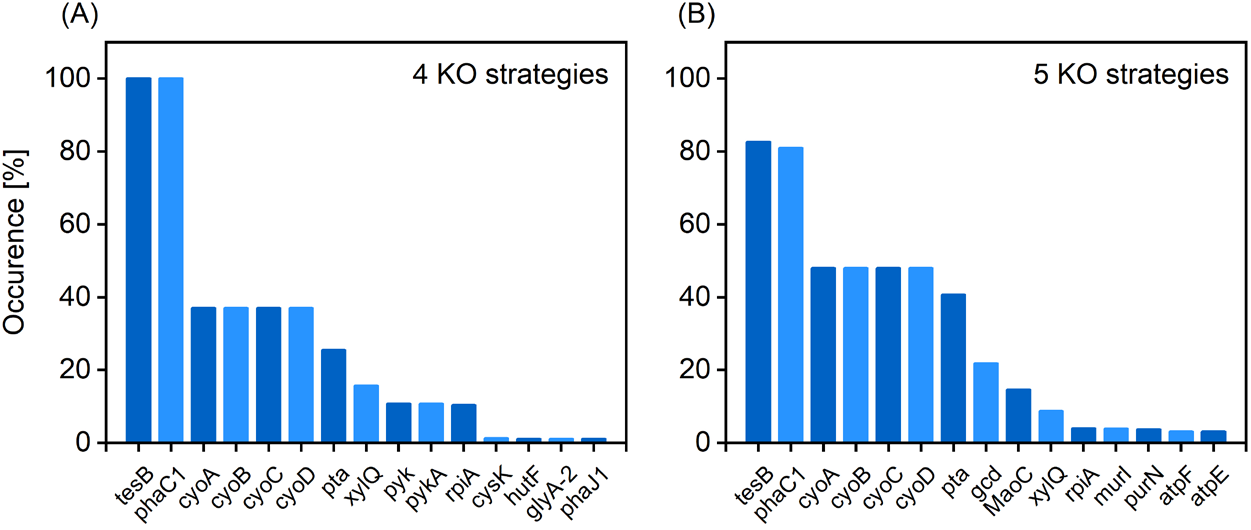
Occurrence distribution of gene knock-out (KO) targets in the best, unique strategies for methyl ketone overproduction determined by the genetic algorithm. (A) and (B) show results for separate genetic algorithm runs searching for quadruple and quintuple KO strategies, respectively. Only strategies that led to an increase in fitness of more than 50 % compared to the reference VLB120 Δ3 pProd were considered.

### 3.6 Methyl ketone production is not limited by precursor drain into polyhydroxyalkanoate synthesis

We initially assessed the impact of individual model-predicted gene deletions in the background of strain VLB120 Δ3 pProd. We first targeted the polyhydroxyalkanoate (PHA) biosynthesis. PHAs are derived from intermediates of the *de novo* fatty acid synthesis and degradation (Figure 1) and are typically produced by *Pseudomonas* as a carbon and energy reserve under noncarbon nutrient-limiting conditions (Brandl et al., 1988). We argued that the upregulated β-oxidation cycle in the engineered *Pseudomonas* strain could induce PHA production under standard growth conditions, which would result in competition for precursors with methyl ketone synthesis. In *P. taiwanensis* VLB120, two PHA synthases are encoded in an operon together with the PHA depolymerase. We deleted the complete *pha* operon (PVLB_02155-PVLB_02165) but did not observe improved methyl ketone production with the resulting strain VLB120 Δ3 *Δpha* pProd in batch shake flask experiments (Figure 2).

### 3.7 Deletion of *tesB* boosts methyl ketone production

The second predicted target, *tesB*, encodes a type II thioesterase, which catalyzes the reversed reaction of the overexpressed FadD. The TesB thioesterase has a substrate specificity towards C12 to C18 (hydroxy)acyl-CoAs (Spencer et al., 1978). In previous metabolic engineering of *E. coli*, this gene was overexpressed to produce enantiopure 3-hydroxybutyrate (Tseng et al., 2009) and to reverse the flux through the β-oxidation (Dellomonaco et al., 2011). However, a knock-out of *tesB* as a strategy to lift the synthesis of fatty acid derivatives has not been reported so far.

Deletion of *tesB* in VLB120 Δ3 pProd increased the total methyl ketone titer by 3-fold to 333 mg L_aq_^-1^ (Figure 2). We further tested the *tesB* gene deletion strain VLB120 Δ3 *ΔtesB* pProd under controlled bioreactor conditions (Figure 5). Under batch conditions, VLB120 Δ3 *ΔtesB* pProd produced 1.3 g L_aq_^-1^ (Figure 5A, Table 2). The increase in methyl ketone titer after carbon source depletion can be explained by a slow, spontaneous decarboxylation of the 3-keto acid. Indeed, we had observed a significant increase in the methyl ketone titer after incubating the fermentation broth of a predecessor strain at 70 °C (Supplementary Figure S4), supporting the hypothesis that the decarboxylation is a limiting step. However, as we observed PHA granules at the end of the batch-phase of a fed-batch fermentation of this strain (not shown), the supply of additional carbon and energy by PHA degradation cannot be excluded. The titer further increased to 6.0 g L_aq_^-1^ in a fed-batch fermentation; productivity and yield also increased (Figure 5B, Table 2). The methyl ketone distribution showed no changes in comparison to the VLB120 Δ3 pProd.

**Figure 5.**
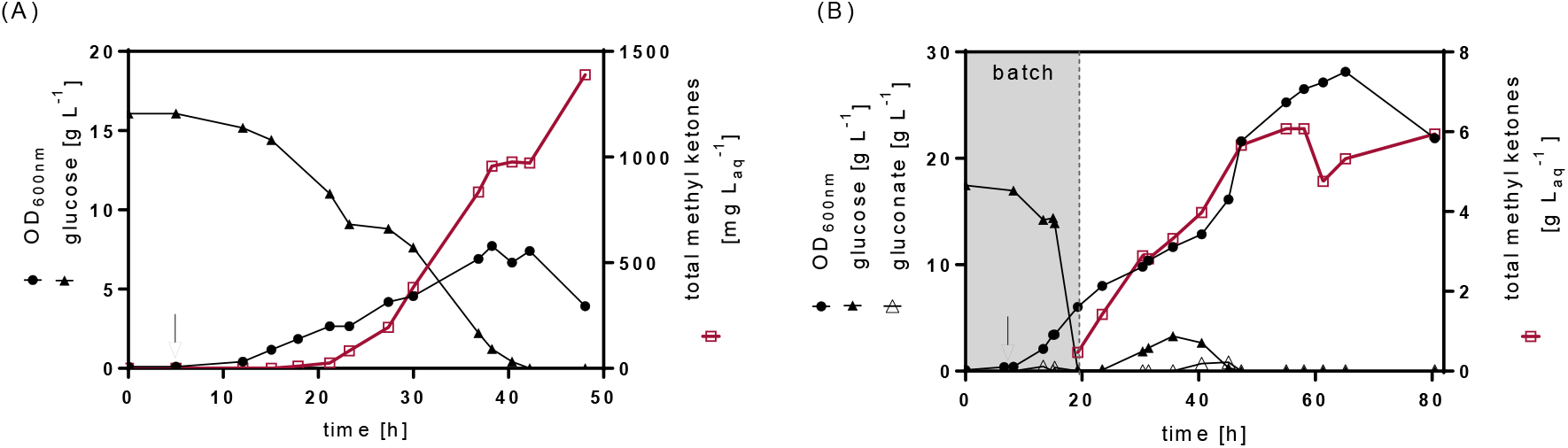
Methyl ketone production and growth during batch (A) and fed-batch (B) cultivation of *P. taiwanensis* VLB120 Δ3 Δ*tesB* pProd in mineral salt medium with an initial concentration of 17 % (v/v) *n*-decane as second organic phase. Shown are the time courses of the optical density and relevant substrates and products. For the reason of clarity, gluconate concentrations, which increased to low levels of 0.1 g L^-1^ in the batch experiment, were excluded from (A). A vertical arrow indicates the time point of induction. For both the batch and fed-batch fermentations, data of one experiment are shown.

### 3.8 Consolidation of individually tested gene deletions further increases production

The two targets predicted by metabolic modeling, *tesB* and *pha*, as well as the gene *fadA2* were deleted in VLB120 Δ3 pProd to test the combined effect of these three knock-outs. When the resulting strain VLB120 Δ6 pProd was cultivated without *in situ* product extraction into an *n*-decane overlay, we observed a white precipitate on the flask bottom after 24 h (Supplementary Figure S5). A GC analysis of the crystal-like precipitate dissolved in *n*-decane verified that the solids contained high amounts of methyl ketones. Only methyl ketone congeners with chain lengths between C13 and C15 were detected, while the C11 ketone probably did not precipitate due to the lower melting point of 15 °C (National Center for Biotechnology Information).

In a 48-h shake flask experiment with *in situ* product extraction, VLB120 Δ6 pProd produced 455 mg L_aq_^-1^ methyl ketones, a 4-fold increase in comparison to VLB120 Δ3 pProd (Figure 2). In fed-batch fermentation, the growth of this consolidated production strain ceased after the batch phase, while the methyl ketone titer significantly increased (Figure 6). VLB120 Δ6 pProd also outperformed the predecessor strains under these cultivation conditions, in which 9.8 g L_aq_^-1^ methyl ketones (corresponding to 69.3 g L in the *n*-decane extraction phase) were produced in less than 80 h (productivity of 132 mg L_aq_^-1^ h^-1^). The product yield increased to 171 mg g_GLc_^-1^, ~53 % of the maximum theoretical yield (Table 2). As with the predecessor strains, C13:1, C15:1, C11, and C13 were the most dominant methyl ketone congeners. The fraction of unsaturated congeners was approx. 60 %, which is significantly higher as reported for *E. coli* (~40 %) and *R. eutropha* (~10 %) (Goh et al., 2012; Müller et al., 2013) but has likewise been observed for engineered *P. putida* (Dong et al., 2019). The abundance of these congeners correlates with the fraction of unsaturated fatty acid moieties in the lipids of these species, reported to be around 70 % for *Pseudomonas* (Rühl et al., 2012), 40 % for *E. coli* (Pramanik and Keasling, 1997) and to vary between 17-30 % for *R. eutropha* (Park et al., 2011).

**Figure 6.**
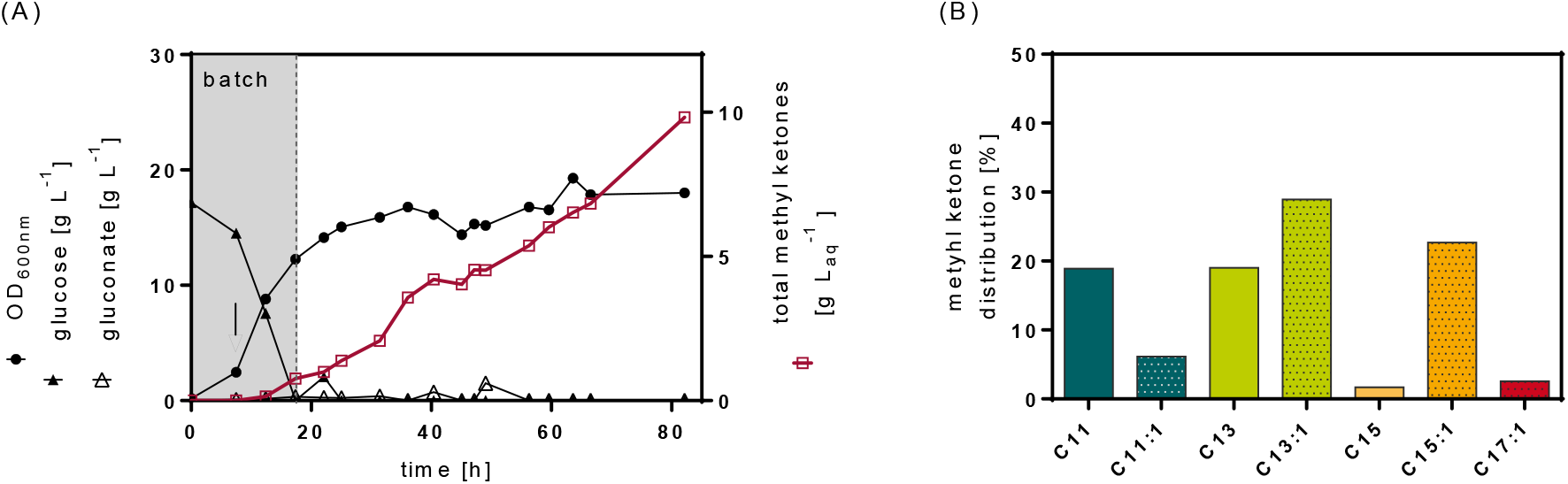
Methyl ketone production in *P. taiwanensis* VLB120 Δ6 pProd in fed-batch cultivation with an initial concentration of 17 % (v/v) *n*-decane as second organic phase. (A) Time course data of optical density and relevant substrates and products. Total methyl ketones represent the sum of all detected methyl ketone congeners. A vertical arrow indicates the time point of induction. (B) Methyl ketone distribution at the end of the fed-batch fermentation.

The proportion of unsaturated fatty acids in *Pseudomonas* is increased to counteract a reduction of membrane fluidity at the lower cultivation temperature of 30 °C (Demendoza and Cronan, 1983; Loffhagen et al., 2004). Accordingly, higher amounts of monounsaturated methyl ketones were observed in *E. coli* when grown at a reduced temperature supporting this relationship (Goh et al., 2012). For the utilization of methyl ketones as biofuel, this elevated fraction of monounsaturated methyl ketones is disadvantageous. While unsaturation can reduce the melting point, it also negatively affects the fuel properties regarding the cetane number (Knothe, 2008). We determined a derived cetane number (DCN) of 54 for the methyl ketone mixture produced by *P. taiwanensis* VLB120 (Supplementary Table S2). Although still in the diesel range, this value is much lower compared to the DCN of approx. 63 calculated for a mixture of fully saturated methyl ketone congeners with the same chain length distribution. This disadvantage of the *Pseudomonas* hosts could be addressed by engineering the specificity of acyl-CoA thioesterases towards saturated fatty acids (Hernandez Lozada et al., 2018) or modulating the fatty acid composition by adapting *Pseudomonas* to growth at higher temperatures.

## 4. Conclusion

By applying rational and model-guided metabolic engineering, *P. taiwanensis* VLB120 was successfully developed for medium-chain length methyl ketones production at high titer and yield surpassing values achieved with alternative hosts. Notably, this high performance was achieved solely by directing the carbon flux through *de novo* fatty acid synthesis towards methyl ketone synthesis. Key genetic modifications necessary for the upregulated β-oxidation to be fully effective were the complete disruption of the β-oxidation cycle and the reverse flux from fatty acyl-CoA esters to free fatty acids. Optimization of the redox metabolism, such as redox cofactor engineering, shown to be important for methyl ketone production in *E. coli*, was not pursued. We argue that the high flexibility of *Pseudomonas* to balance redox cofactors (Kohlstedt and Wittmann, 2019; Nies et al., 2020; Nikel et al., 2016) renders such elaborate engineering efforts superfluous. Moreover, the performed *in silico* analysis showed that the capacity of the strain for methyl ketone production has not been fully exhausted and can be increased further by metabolic engineering. The early evaluation of cultivation strategies also showed the impact of the fermentation settings on the strain performance with nitrogen-limited, carbon-excess conditions benefitting a high flux towards methyl ketone synthesis. We believe that the full potential of *P. taiwanensis* VLB120 remained untapped in the initial, unoptimized fed-batch experiments as the rather low glucose feed rate prevented utilization of *P. taiwanensis* VLB120’s high redox cofactor regeneration capacity for maximal methyl ketone synthesis. Future fermentation process optimization will further leverage the strain’s metabolic capability for redox cofactor supply and conversion of non-edible renewable resources or waste streams such as lignocellulosic biomass or glycerol.

## Abbreviations

aq: aqueous phase
CDW: cell dry weight
DO: dissolved oxygen
GC: gas chromatography
GLC: glucose
HPLC: high-performance liquid chromatography
MiMBl: Minimization of Metabolites Balances
MK: methyl ketone
MS: mass spectrometry
OD_600nm_: optical density determined at 600 nm
org: organic phase
(p)FBA: (parsimonious) flux balance analysis

## Acknowledgment

We are grateful to Andrea König (AVT, RWTH Aachen University, DE) for the prediction of fuel properties.

## Supplementary Material

**Supplementary Figure S1.**
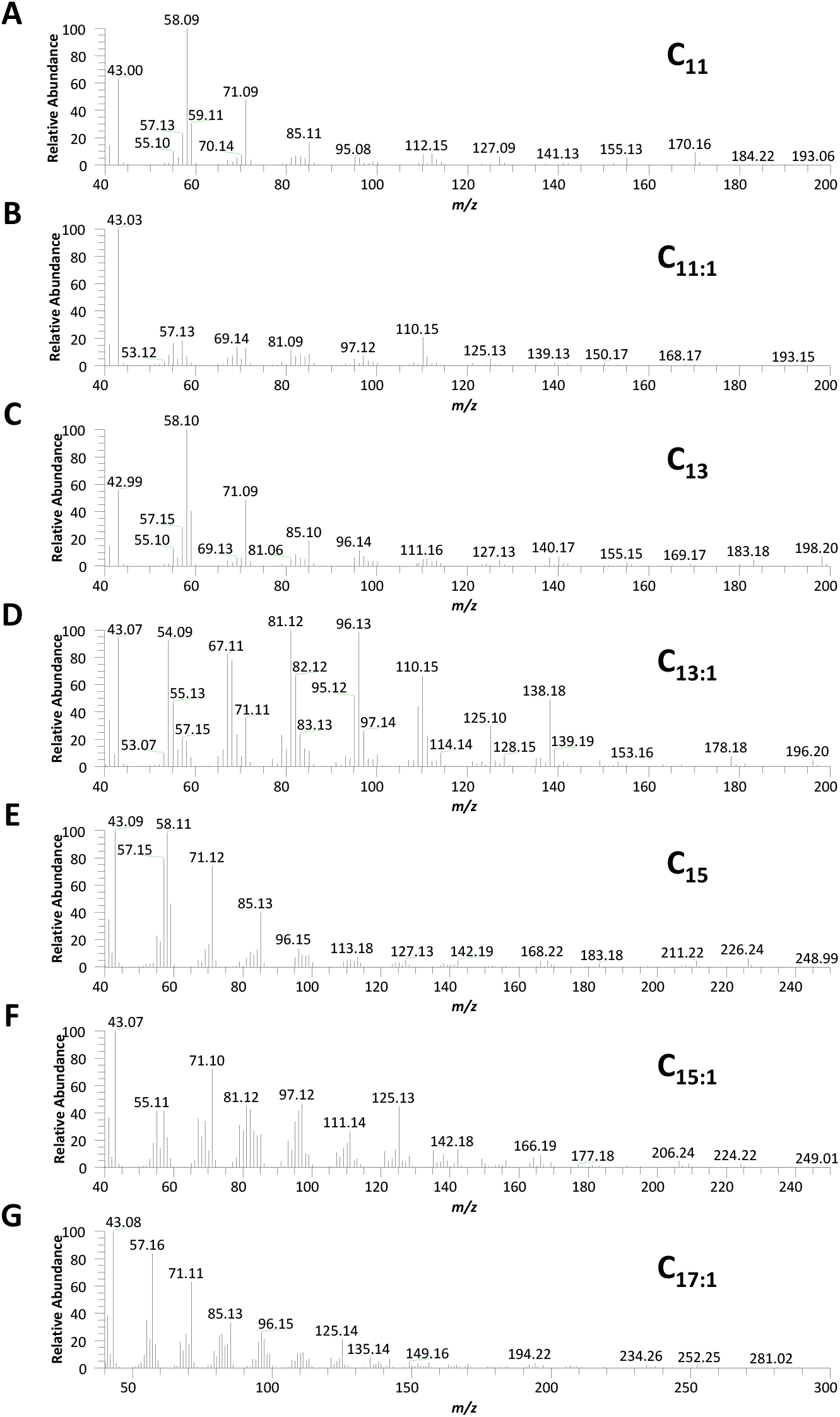
Mass spectra of methyl ketones generated by VLB120 Δ6 pProd at 66 h. (A) C_11_, (B) C_11:1_, (C) C_13_, (D) C_13:1_, (E) C_15_, (F) C_15:1_, and (G) C_17:1_. All saturated methyl ketones were identified with authentic standards. The monounsaturated methyl ketones were tentatively identified by comparison with spectra shown in (Goh et al., 2012).

**Supplementary Figure S2.**
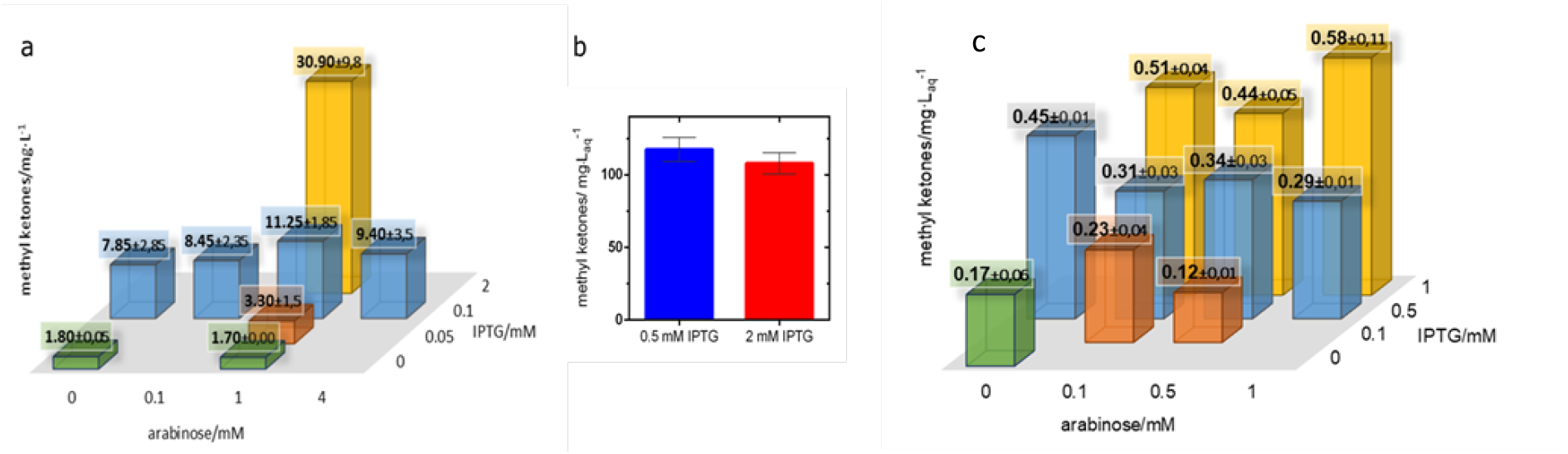
Determination of optimal inducer concentrations for methyl ketone production. The inducers IPTG and arabinose were added to the shake flask cultures 4.5 h after inoculation. The methyl ketone titers shown are the mean of duplicate experiments with the corresponding standard deviation and were measured after 24 h of cultivation. a) VLB120 Δ3 pProd was grown in 100 mL shake flasks without *n*-decane overlay. The overall titer of detected methyl ketone congeners (C11, C13, C15) is shown. b) VLB120 Δ3 pProd was grown in 250 mL shake flasks with *n*-decane overlay and induced with 1 mM arabinose and either 0.5 or 2 mM IPTG. The overall titer of detected methyl ketone congeners (C11, C13, C13:1, C15, C15:1) related to the aqueous phase is shown. c) VLB120 Δ3 *att*Tn7::MK was grown in 100 mL shake flasks with a *n*-decane overlay. The overall titer of detected methyl ketones congeners (C11, C13, C13:1, C15, C15:1) related to the aqueous phase is shown.

**Supplementary Figure S3.**
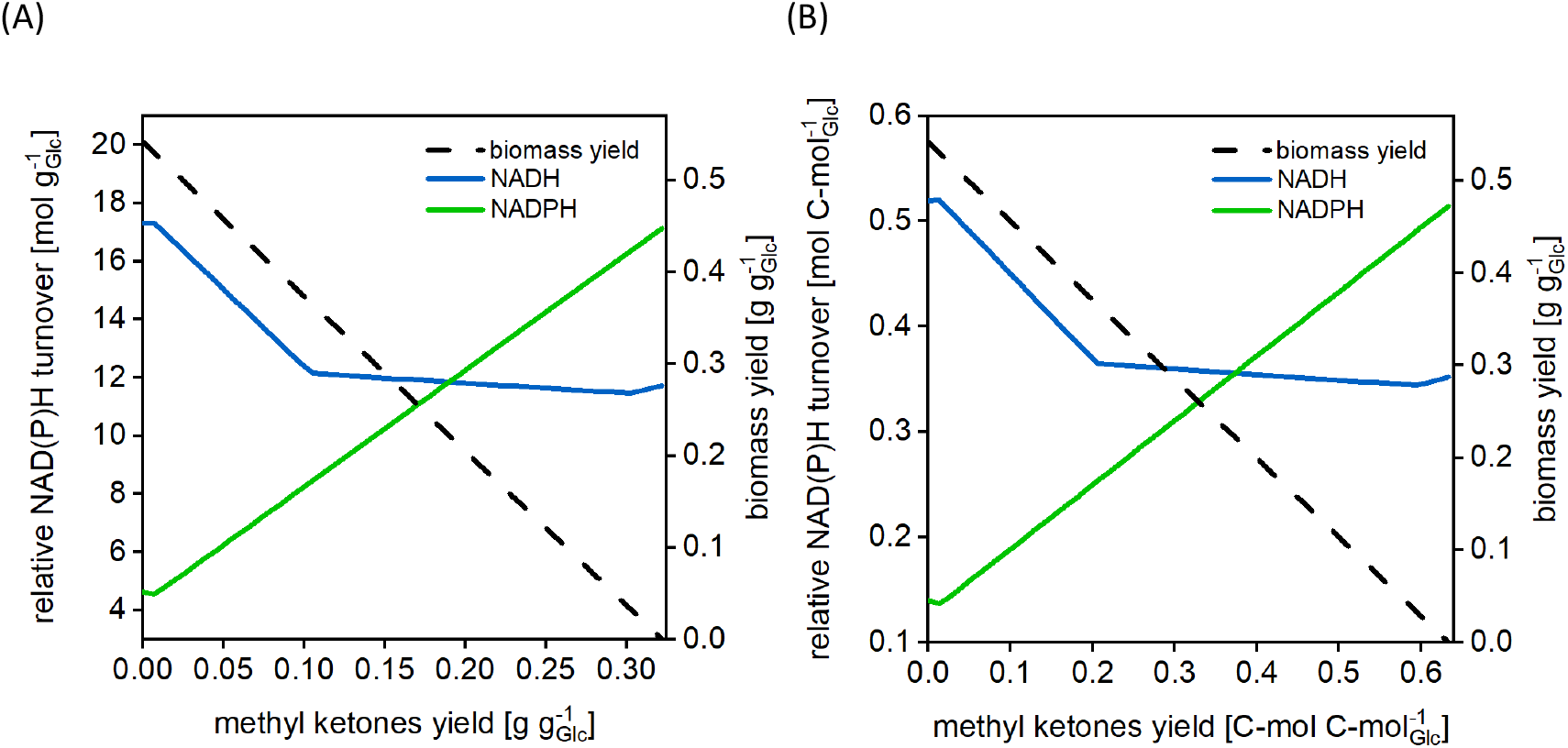
Simulated NAD(P)H turnover and biomass yield as a function of a given methyl ketone yield. (A) mass related, (B) on C-mol basis. The NAD(P)H turnover values and biomass yields were derived from growth-optimal flux distributions at a fixed methyl ketone production rates. Biomass yields were computed by dividing the growth rate by the glucose uptake rate. The NAD(P)H turnover depicts the sum of fluxes of all reactions that produce NAD(P)H divided by the glucose uptake rate. All NAD(P)H producing fluxes were multiplied with the stoichiometric coefficient of NAD(P)H for the respective reaction. Note that for constraint-based modeling approaches, as applied here, metabolite concentrations are assumed to be constant. Thus, summation of all NAD(P)H *consuming* fluxes yields the same NAD(P)H turnover values. The *i*JN1411 model, including the methyl ketone production pathway, was used for all simulations. The methyl ketone distribution was set as depicted by the experimental results for VLB120 Δ3 pProd (cf. Results, section 3.3).

**Supplementary Figure S4.**
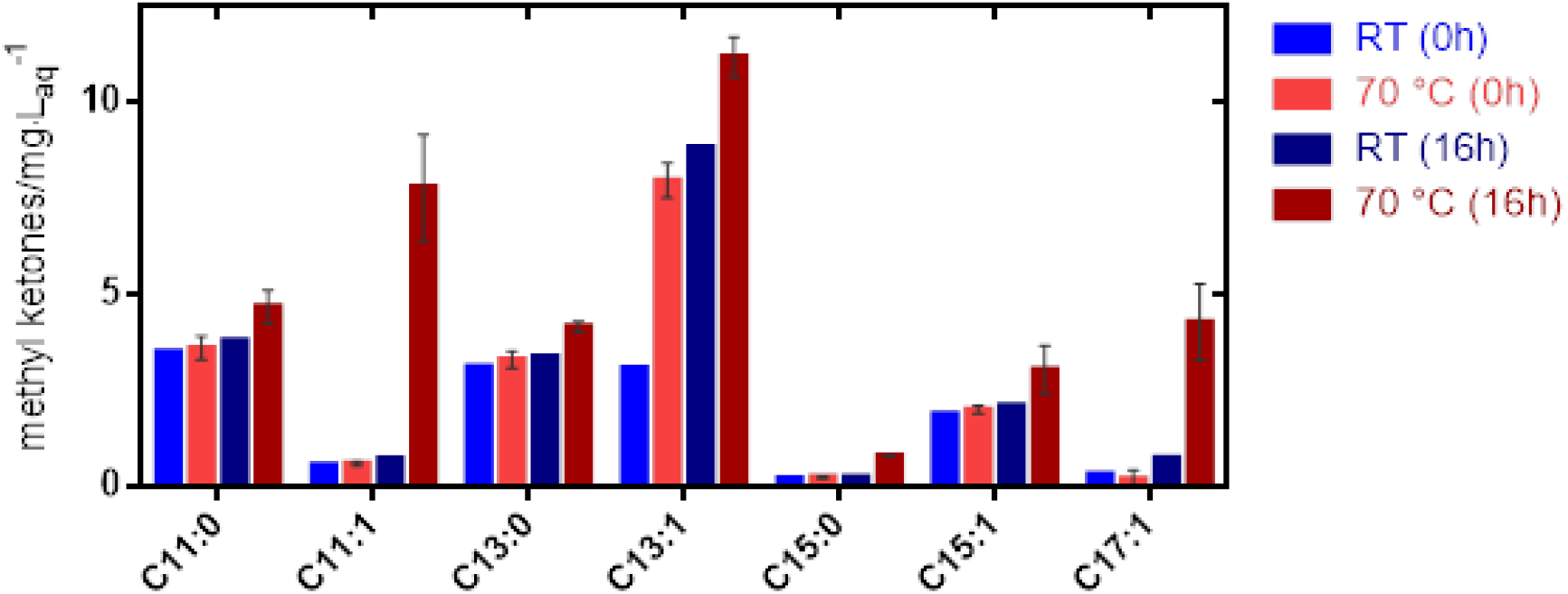
Enhanced decarboxylation of 3-ketoacids by incubation of culture broth at 70 °C for 16 h increases methyl ketones titers. Three samples from a culture of VLB120 Δ3 pProd were taken; two were incubated at 70 °C while the third sample was left at room temperature (RT). The concentrations of saturated (C11, C13, C15) and monounsaturated (C11:1, C13:1, C15:1, C17:1) methyl ketone congeners were determined before (0h) and after the incubation period (16h). Error bars represent the standard deviation of the duplicate experiment at 70 °C.

**Supplementary Figure S5.**
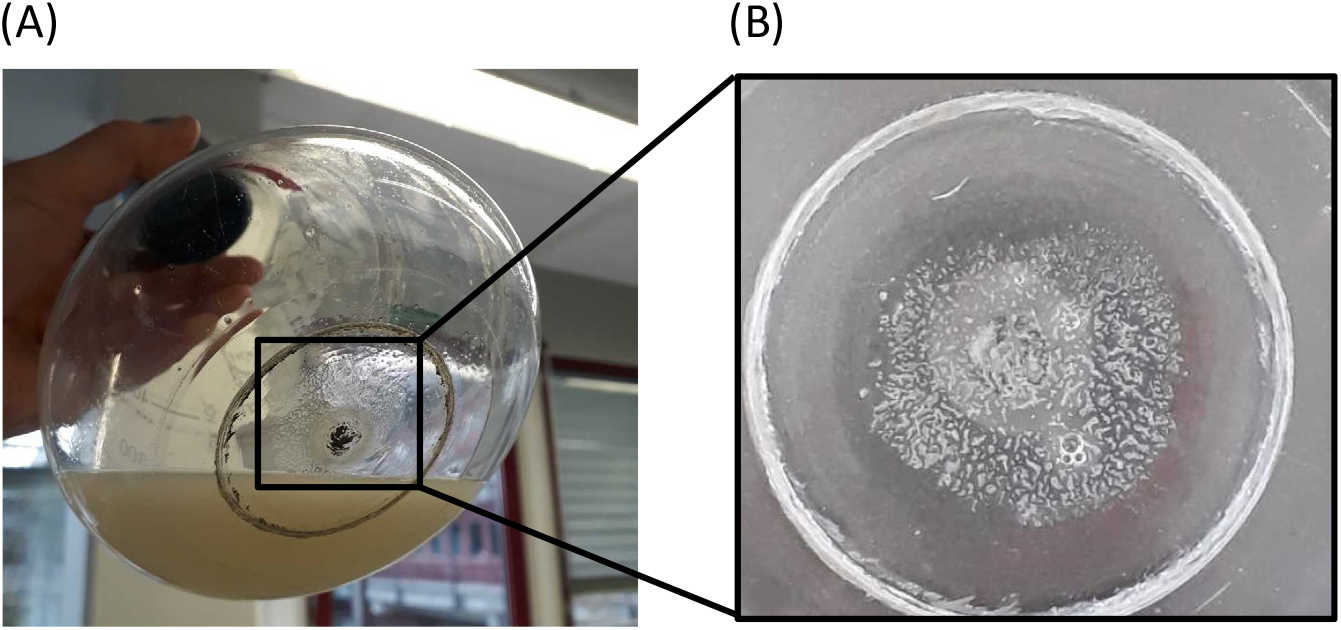
Cultivation of *P. taiwanensis* VLB120 Δ6 pProd in mineral salt medium without *n*-decane as an organic phase. White crystal-like precipitation on the flask bottom after 24 h cultivation. (B) View of the flask bottom from above.

**Supplementary Table S1.**
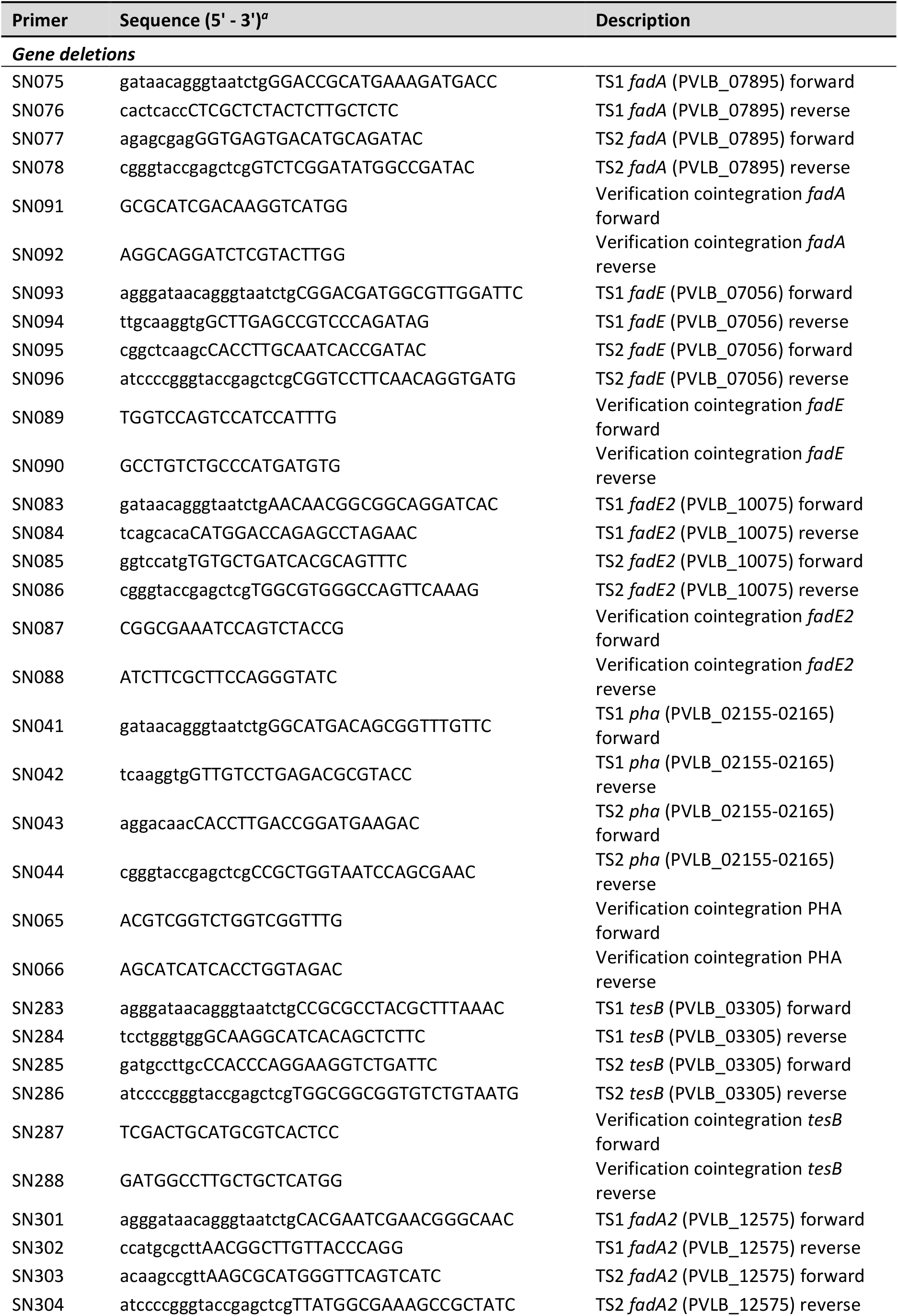
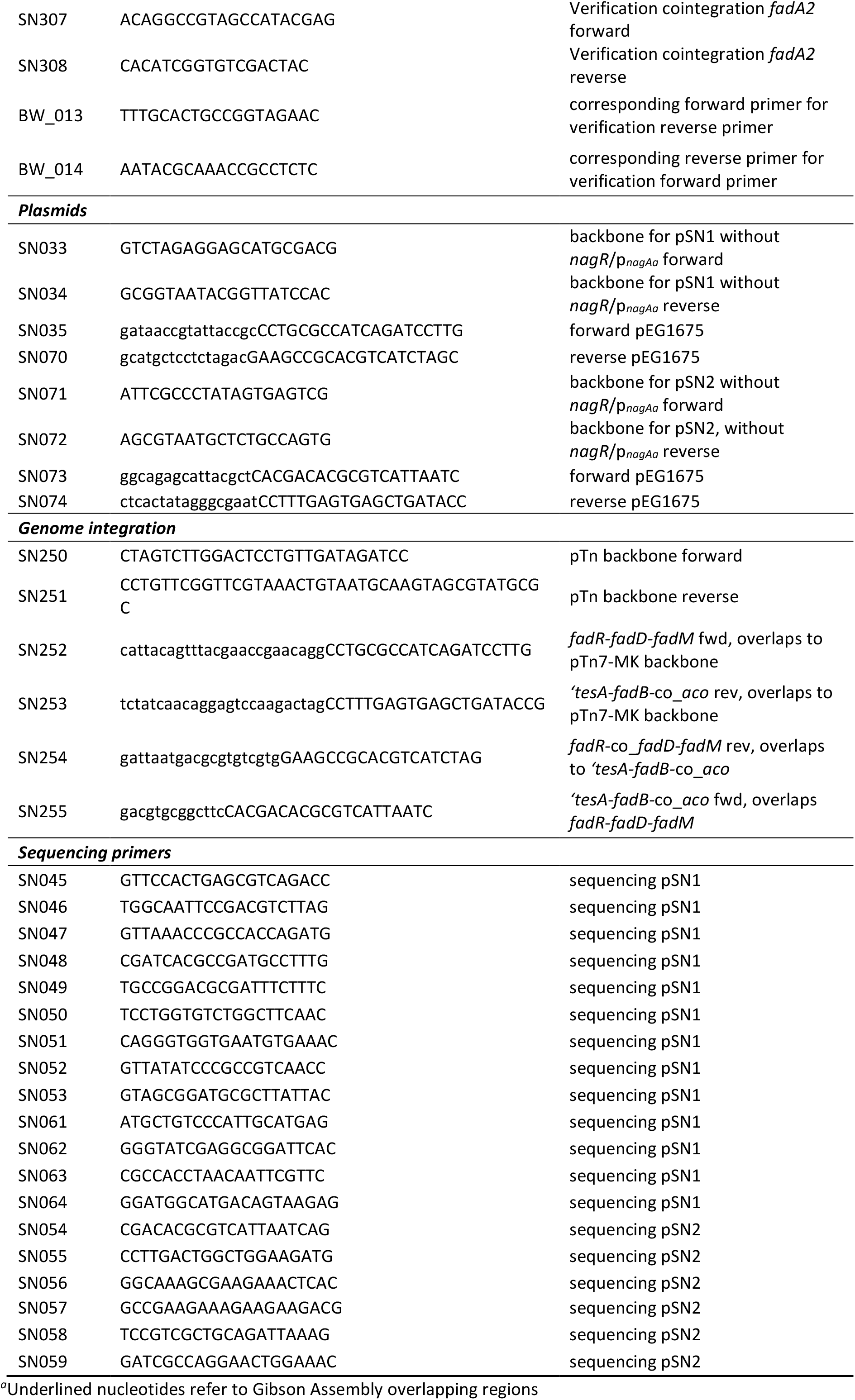
Primers used in this study

**Supplementary Table S2.** Fuel properties of the produced methyl ketone (MK) mixture

| MK species | C11 | C11:1 | C13 | C13:1 | C15 | C15:1 | C17:1 |
| --- | --- | --- | --- | --- | --- | --- | --- |
| **Mol fraction (VLB120 Δ3 pProd)**<br>[mol/mol] | 0.26 | ND | 0.20 | 0.31 | 0.01 | 0.19 | 0.02 |
| **Derived cetane number (DCN)** | 51.8 | 39.7 | 62.3 | 50.1 | 76.6 | 60.4 | 72.9 |
| **Boiling point [°C]** | 224.0 | 221.7 | 255.8 | 253.6 | 282.5 | 279.8 | 306.5 |

The derived cetane number (DCN) of a methyl ketone (MK) blend was determined by assuming linear additivity (Dahmen and Marquardt, 2017; Knop et al., 2014) and using the following equation, in which $x_i$ and $DCN_i$ designate the mole fraction and the DCN of component $i$ , respectively (Dahmen and Marquardt, 2017; Knop et al., 2014):

